# Social media at metabolic meetings: who is tweeting what and for whom?

**DOI:** 10.1101/2022.08.16.504099

**Authors:** James Nurse

## Abstract

**BACKGROUND:** The social media site Twitter has been widely embraced in medical circles for its ability to connect individuals and support rapid information sharing. Critics say that the messages shared may not accurately reflect what was said and that sharing meeting content could devalue conferences themselves. It is unclear how it is used at SSIEM and what value it may bring.

**METHODS:** Twitter’s tweetdeck software was used to find all tweets containing the conference ‘hashtag’ #SSIEM2018. All tweets were reviewed to identify the author, see what had been shared and count replies, likes and retweets. Authors were grouped by professional background and tweet content was broken down by type of material shared and theme.

**RESULTS:** 122 relevant tweets were sent during the fortnight at the beginning of September 2018, creating over 400,000 impressions. There were a further 73 replies with approximately 13 engagements (likes, replies or retweets) per tweet. 36 people wrote tweets (rate: 3.4 per person [1-33]). One quarter of the tweets shared poster content and over one third of tweets related to Phenylketonuria materials. 50 of the tweets were produced by just two accounts, both intended to provide information to patients and their families.

**DISCUSSION:** Tweets where no hashtag was used cannot be identified and restrictions within Twitter prevent certain analyses on tweet data greater than 30 days old. However, Twitter uptake within metabolic medicine is significantly behind other specialities where conference tweets can exceed 20,000. Information shared is typically intended for patients rather than other health professionals; this suggests a different uptake to more mainstream specialities. Presenting teams should be aware that their work may be received directly by patients and families and consider how best to present their messages for all who may receive them.

## BACKGROUND

Social media refers to websites and applications designed to allow the sharing of user-generated content, such as text, photos or video. Twitter is one such application, allowing users to upload ‘tweets’ of up to 280 characters that are seen by their ‘followers’ and anyone who chooses to look for them. ‘Tweets’ can be labelled with ‘hashtags’ allowing them to be grouped around a particular theme or event; this hashtag can then be found by other users.

Twitter use is rife; 500 million tweets are sent a day or 5,787 tweets every second. Although the total number of registered users is falling, 326 million people use Twitter every month including a quarter of US adults. Recall from Twitter is felt to be better than other online sites and many Twitter users get their news on Twitter rather than other online sites^1^.

Twitter has been rapidly adopted by the medical community and is used for education, networking and sharing research^2 3^. It is the favoured social network for communication before, during and after conferences^4^ and most meetings will now have a designated hashtag and encourage attendees to ‘get involved’. The International Conference on Emergency Medicine in 2012 saw over 4500 tweets from 400 people tweeting although only 1/3 were at the meeting itself^5^. In 2014 the Social Media and Critical Care Conference (SMACCGold) had 23000 tweets over 4 days with over 39 million impressions^6^.

Although widely used in some fields of medicine, other specialities have been less keen to adopt Social Media and Twitter. Within the Inherited Metabolic Disease (IMD) community there are some very engaged groups, notably in the UK the metabolic dieticians at Birmingham Children’s Hospital share daily advice on low protein foods and Phenylketonuria (PKU) related papers. Clinicians working in IMD are less obvious online. The reason for the absence of metabolic clinicians is unclear although the public response to cases such as that of Charlie Gard^7^ and Alfie Evans may explain the low profile. However, a failure to engage does not prevent the ever pervasive presence of social media and it is important o consider what role it may have to play in IMD.

## OBJECTIVES

To analyse Twitter usage around a metabolic meeting to identify the content shared, the role of the person sharing content and the number of users reached.

## METHODS

Twitter’s tweetdeck software was used to find all tweets containing the conference ‘hashtag’ #SSIEM2018. Tweets were reviewed to identify the author, see what had been shared and count replies, likes and retweets. Authors were grouped by professional background and tweet content was broken down by type of material shared and theme.

## RESULTS

122 tweets containing the #SSIEM2018 tag were sent during the fortnight at the beginning of September 2018, with an estimated reach of 400,000 (no. of tweets * no. of followers).

**Fig 1:**
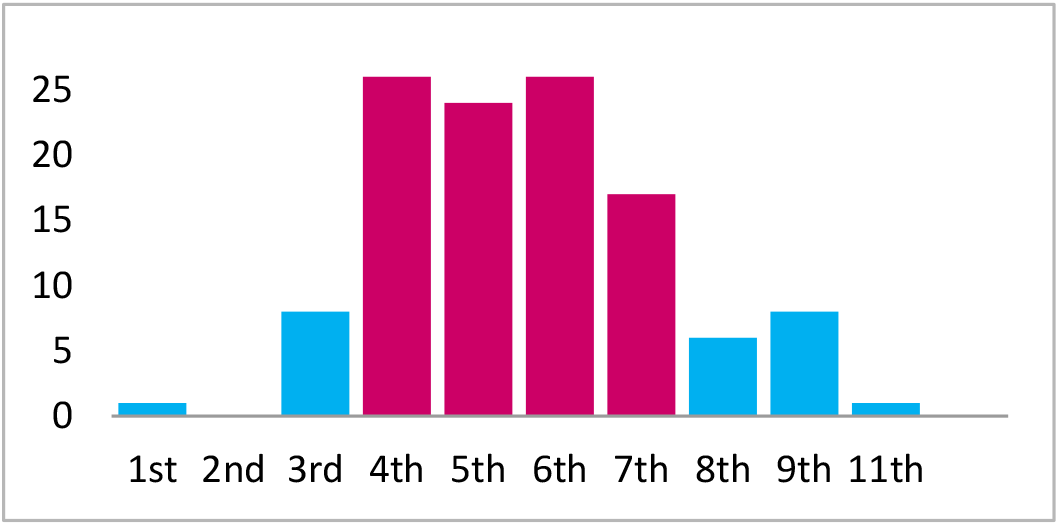
No of tweets per day using #SSIEM2018 in September 2018

There were a further 73 replies to these tweets and, review of specific twitter data for four of the users identified a small number of tweets that did not use the conference hashtag. In total there were approximately 200 meeting tweets.

The 122 primary tweets had approximately 13 engagements (likes, replies or retweets) per tweet; does not include ‘link clicks’ or ‘picture views’ that twitter also counts as engagements.

36 people wrote tweets (rate: 3.4 per person (1-33)). 50 of the tweets were produced by just two accounts, both intended to provide information to patients and their families. Patient groups and dieticians wrote over half of the tweets sent.

**Fig 2:**
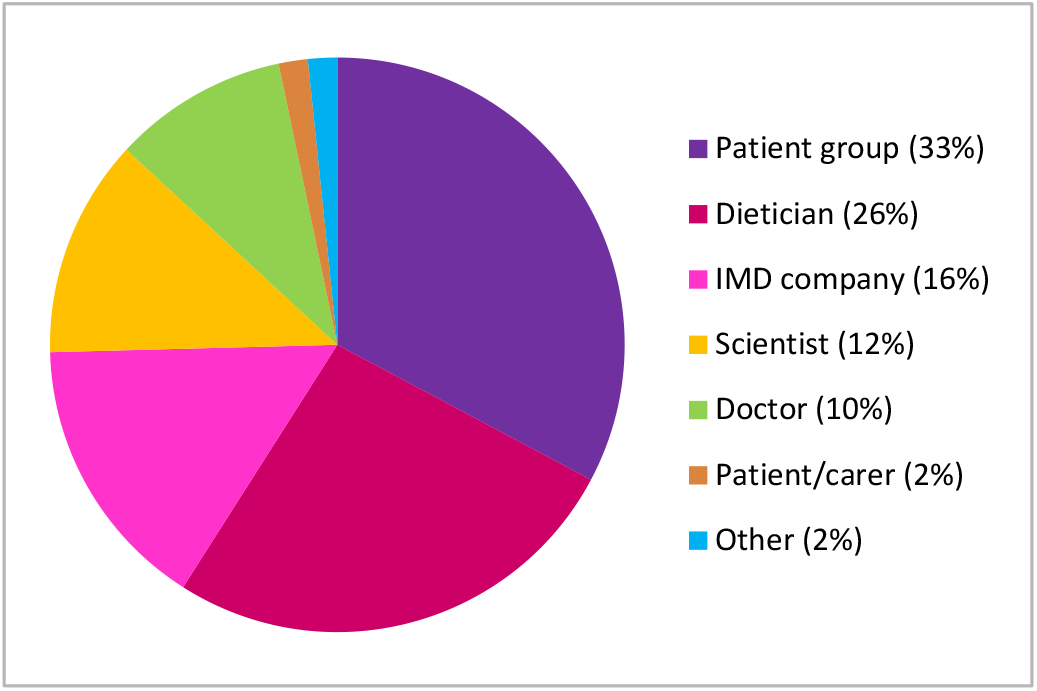
Make-up of users for 122 tweets containing #SSIEM2018 tag

One quarter of the tweets shared poster content and over one third of tweets related to Phenylketonuria. Almost half of the tweets contained non-specific meeting or attendee details without any clinical/scientific content.

**Fig 3:**
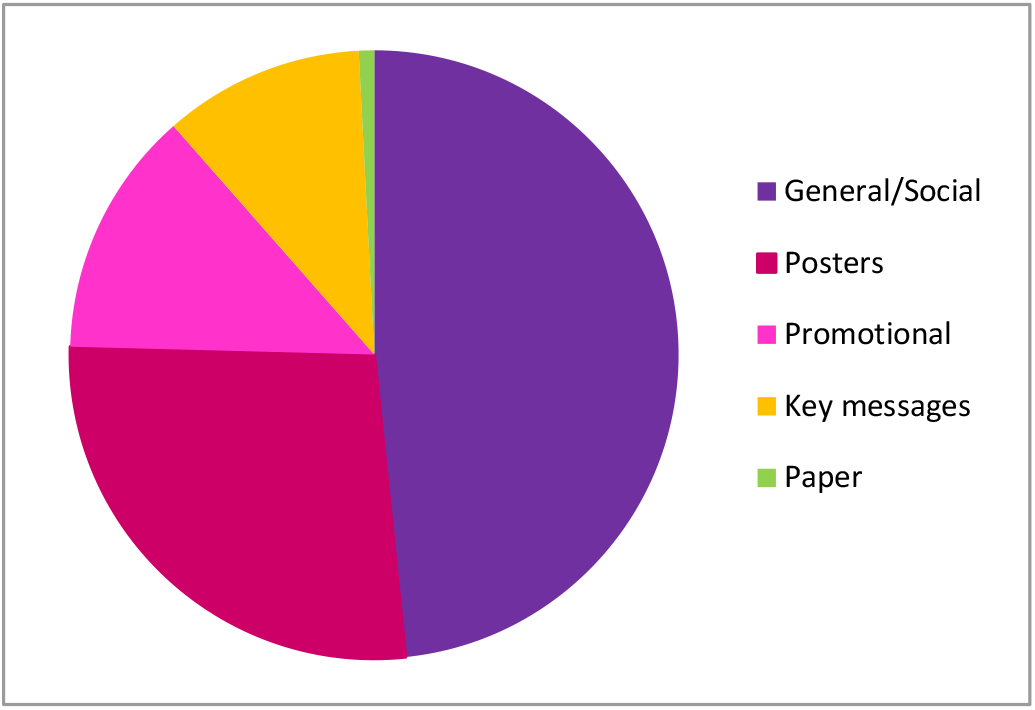
Content of tweets containing #SSIEM2018 tag

## DISCUSSION

The approximately 200 tweets identified as related to the 2018 SSIEM meeting is significantly behind the activity seen for meetings in other specialities. It is unclear why adoption is low within IEM; notably clinicians are very poorly represented. It may be that a critical mass needs to be reached to generate further uptake.

4 of the 36 users, including the two most prolific tweeters (@NSPKU and @metabolicsBCH), shared twitter analytic data for the dates over which their SSIEM tweets occurred. This enabled identification of some unlabelled tweets and also gave accurate impression and engagement data.

This extended data for 59 of the primary tweets showed average impressions of 1100 users with 54 engagements per tweet – this higher figure includes users clicking on the tweet to see embedded photos or viewing a user’s profile.

Of the five tweets with the most impressions, two shared scientific detail from posters and three contained no clinical information at all; all five were PKU related.

Poster information that was shared often involved pictures with the author suggesting some consent for sharing; consent cannot be presumed and may not be in keeping with the policies of the meeting’s organisers.

This work is limited by the inability to identify tweets where a relevant hashtag was not used. Twitter also restricts the availability of analytic data around hashtags to the last 30 days; this limits the opportunity to generate accurate impressions data for all tweets and the #SSIEM2018 tag itself.

@NSPKU and @metabolicsBCH largely share information for followers interested in PKU news. Their activity at the meeting therefore biases the information shared. However, it does illustrate how tweets from a meeting can be shared for non-professionals and the subsequent interest in these messages.

## CONCLUSION

Social media uptake in metabolic medicine is low. Roland and Brazil make an excellent case for its role in continuing professional development, information sharing and education^8^; not using it may be to the profession’s detriment. The messages that speakers share are typically correctly reported, although this may not always be the case^9^. It is unclear how much harm may arise from the misrepresentation of shared information.

Increasingly Twitter is used as a way to speak directly to a user base, as demonstrated by the current President of the United States^10^. This can present an excellent opportunity to rapidly disseminate information and researchers may be keen to share news with their intended patient base; families are keen to hear news in an evolving field of medicine where conditions may have no current treatment.

It is clear that Twitter is there for those who tweet and, in Inborn Errors of Metabolism that remains the minority. However, teams should be aware that their work may be received directly by patients and families and they may wish to consider this when they decide how best to present their findings.

## GLOSSARY

Engagement: The number of interactions people have with your content (i.e.: likes, comments, link clicks, profile clicks, retweets, etc.)^11^
Hashtag: The # symbol, called a hashtag, is used to mark keywords or topics in a Tweet. It was created organically by Twitter users as a way to categorize messages^12^.
Impressions: the number of times your content is displayed to users^11^.
Reach: the number of people who could see your content^11^.
Reply: A Tweet posted in reply to another user’s message, usually posted by clicking the “reply” button next to their Tweet in your timeline. Always begins with @username^12^.
Retweet: (verb) To retweet, retweeting, retweeted. The act of forwarding another user’s Tweet to all of your followers. (noun): A Tweet by another user, forwarded to you by someone you follow. Often used to spread news or share valuable findings on Twitter^12^.
Social Media: Interactive computer-mediated technologies that facilitate the creation and sharing of information, ideas, career interests and other forms of expression via virtual communities and networks. Typically built on web 2.0, featuring user generated content shared from specific user profiles and commonly designed to support the development of social networks.
Twitter: a microblogging and social networking service on which users post and interact with messages known as “tweets”.

